# Focal cholesterol depletion during Fcγ receptor-mediated phagocytosis contributes to Lyn kinase regulation

**DOI:** 10.1101/388637

**Authors:** Stella M. Lu, Allen Volchuk, Gregory D. Fairn

**Author notes:** Corresponding author: Gregory D. Fairn. Address correspondence to: Gregory D. Fairn, Keenan Research Centre for Biomedical Science, Office #614, LKSKI Building, St. Michael’s Hospital, 209 Victoria Street, Toronto, ON M5B1W8, CANADA.

## Abstract

Cholesterol-rich nanodomains, historically referred to as lipid rafts, have previously been reported to be critical for proper Fcγ Receptor and Lyn kinase signaling during phagocytosis. Throughout the initial stages of phagocytosis, the nascent phagosome is actively remodeled by localized lipid metabolism and exocytosis. However, to date, little is known about the dynamics of cholesterol during this stage of particle engulfment. Using a genetically-encoded biosensor for cholesterol, we find that cholesterol is depleted from the nascent phagosome prior to sealing. Additionally, protein markers of both cholesterol-rich and cholesterol-poor nanodomains also clear from the site of phagocytosis arguing against the selective depletion of specific membrane domains. Consistent with previous studies we find that exocytosis contributes to the remodeling of the nascent phagosome. The displacement of cholesterol from the forming phagosome was paralleled by Lyn kinase helping to explain the reduction of phosphotyrosine signal in the nascent phagosome. This diminution of cholesterol and Lyn from the base of the cup may aid in the processivity of the phagocytic signal during pseudopod extension, and provide an unappreciated mechanism by which Lyn kinase signaling is regulated during phagocytosis.

Summary Statement: Localized exocytosis dilutes cholesterol from the phagocytic cup leading to the displacement of Lyn kinase and an attenuation of signaling.

## Introduction

Phagocytosis, the ingestion of particles >0.5 μm is critical for the elimination of apoptotic cells and potentially harmful microbes. This process is used extensively by professional phagocytes including macrophage and neutrophils as part of the first line of defense of the immune system. Phagocytosis is first initiated by the binding of ligands to surface receptors that transduce signals leading to extensive remodeling of the actin cytoskeleton and the plasma membrane (PM). This leads to the engulfment of particles into a vacuole termed the nascent phagosome. This organelle then matures into highly degradative phagolysosomes following a series of fission and fusion reactions with endosomes and lysosomes.

Of the phagocytotic receptors, the Fcγ receptors (FcγR) that recognize the Fc portion of immunoglobulins are the best understood. Activation of the FcγR leads to the activation of the Src family kinase Lyn which in turn phosphorylates Syk kinase resulting in the extensive actin and membrane dynamics. Previous results have demonstrated that FcγR transitions from detergent-soluble fractions to detergent-insoluble fractions following receptor activation^1^. Historically, these detergent-resistant membranes have been described as “lipid rafts” although this designation has fallen out of favor with many cell biologists^2^. Nevertheless, treatments that either remove or oxidize cholesterol are known to impact the functionality of the FcγR^1,3^.

The roles for cytoskeletal rearrangements and phosphoinositide metabolism during the formation of the nascent phagosome have been well documented. Comparatively, little is known about the dynamics of cholesterol during the formation and maturation of the phagosome. This dearth of knowledge is due, at least in part, to the inability to visualize cholesterol in real time. Recently, we developed fluorescently labeled biosensors based on Domain 4 (D4) of the theta toxin from *Clostridium perfringens* that can be used to monitor cholesterol in living cells^4^. To detect cholesterol in the outer leaflet of the PM a recombinant GFP-tagged version of D4 is added to the extracellular medium^4^. To detect cytosolic leaflet cholesterol, we use a mutant D4 with a lower threshold requirement to bind to cholesterol-containing membranes. Using these probes together with the fluorescent cholesterol binding macrolide, filipin, we have investigated the dynamics of cholesterol during the early stages of phagocytosis.

## Results and Discussion

### D4H binding to the PM is cholesterol-dependent and displaced during phagocytosis

RAW macrophages were transfected with TdTomato-D4H, hereafter referred to as D4H, to monitor the relative abundance of cholesterol in the cytosolic leaflet of the PM. The probe localized to the PM in resting cells (**Figure 1A**) and relocalized to the cytoplasm following the extraction of cholesterol using methyl-β-cyclodextrin (mβCD). The cytoplasmic pool of D4H was also able to detect the re-addition of cholesterol and decorated the PM following the reintroduction of its ligand. These results indicate that D4H is responsive to changes in the plasmalemmal cholesterol.

**Figure 1.**
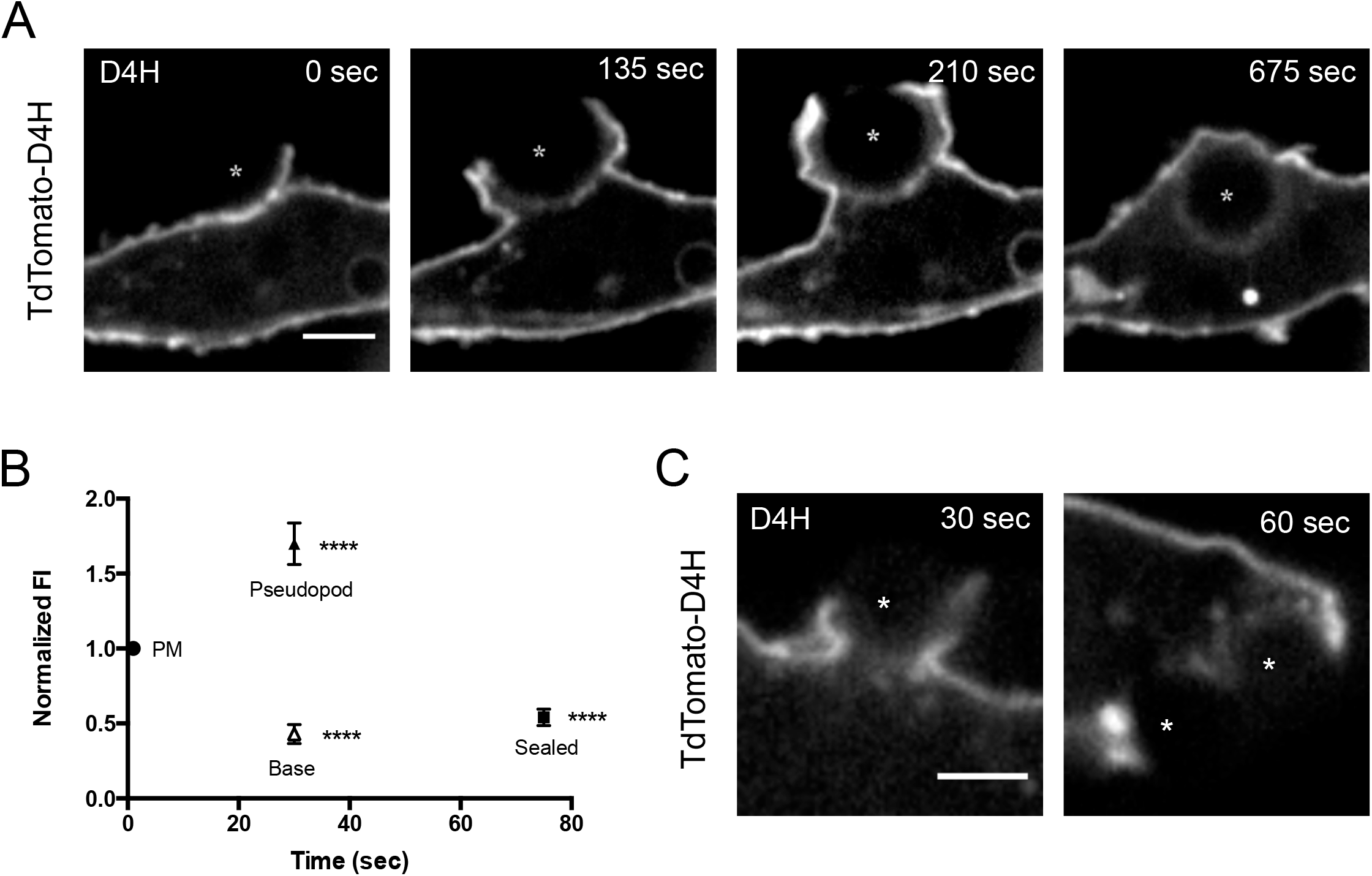
Time-lapse movie of RAW macrophage transfected with TdTomato-D4H and given IgG opsonized polystyrene beads. Asterisks denote phagocytic target and inset shows enlarged area of cell. Scale bar, 10 μm. **(C)** Quantification of the fluorescence intensity of the phagosome during phagocytosis over time. Fluorescence intensity of the PM (unengaged region of the cell), pseudopod (tip of the pseudopod), base (base of the phagocytic cup), and phagosome (phagosomal membrane once phagosome forms) were measured using the equation, (area of interest-background)-(cytosol-background)/(PM-cytosol) at the indicated time points. Fluorescence intensity measurements were normalized to the PM at time 0. At least 20 cells were assessed per condition. A Student’s two-tailed unpaired t-test was performed. ****P< 0.0001 **(D)** COS-IIA cells transfected with TdTomato-D4H and given IgG opsonized polystyrene beads for the indicated time points. Asterisks denote phagocytic target. Scale bar, 10 μm.

To examine the dynamics of cholesterol during phagosome formation we transfected RAW macrophage cells with D4H and challenged the cells with IgG opsonized particles. During phagocytosis, we found that D4H was depleted from the base of the phagocytic cup while being partly enriched towards the front of the migrating pseudopods (**Figure 1B, C**). Once internalized there was an apparent decrease in the intensity of the D4H in the nascent phagosome compared to the unengaged PM. This is consistent with reports that demonstrated that early −10 min old phagosomes-are depleted of cholesterol relative to the PM^5^. We extended this observation to the normally non-phagocytic COS-1 cells. In this experiment, COS-1 cells rendered phagocytic by stably expressing the FcγRIIa, hereafter referred to as COS-IIA, were transiently transfected with D4H and challenged with IgG-opsonized particles^6^. As shown in **Figure 1D**, COS-IIA cells displayed similar D4H dynamics during phagocytosis, with D4H present in the PM and enriched in pseudopods and reduced at the base of the phagocytic cup and nascent phagosome. Collectively, these results demonstrate that cholesterol is locally depleted during phagocytosis and that the mechanism of this diminution is inherent to both macrophage and engineered phagocytic cells.

### Depletion of exofacial and cytosolic cholesterol during phagocytosis

The apparent loss of cholesterol from the inner leaflet could be the result of flopping to the exofacial leaflet. Indeed, previous studies have demonstrated that the ABC transporter ABCA7 is required to support phagocytosis^7^. The mammalian ABCA7 shares high sequence similarity to the *Caenorhabditis elegans* CED-7 gene which was also shown to be required for phagocytosis of apoptotic cells^7^. We examined the distribution of both a GFP-tagged recombinant version of D4 and filipin during phagocytosis. Macrophages were transfected with D4H to visualize inner leaflet cholesterol and incubated with recombinant His6x-GFP-D4 to label exofacial cholesterol (His-D4). During the course of phagocytosis both cholesterol probes displayed a similar localization suggesting that flopping to the outer leaflet was not accounting for the reduction in D4H signal at the base of the cup and nascent phagosome (**Figure 2A, C**). There is no difference in the fluorescence intensities of D4H or His-D4 labeled cells (**Figure 2C**). While not particularly useful for live-cell imaging, filipin has been widely used to visualize cholesterol in healthy cells and cells with lipid storage disorders such as Niemann-Pick Type C^8–10^. Staining RAW cells with filipin revealed that both D4H and filipin signals were diminished from the base of the phagocytic cup during phagocytosis (**Figure 2A, C**). Collectively, our results using the D4-derived porbes and filipin, together with previous biochemical determinations demonstrate that cholesterol is depleted specifically at the base of the phagocytic cup during the formation of the nascent phagosome.

**Figure 2.**
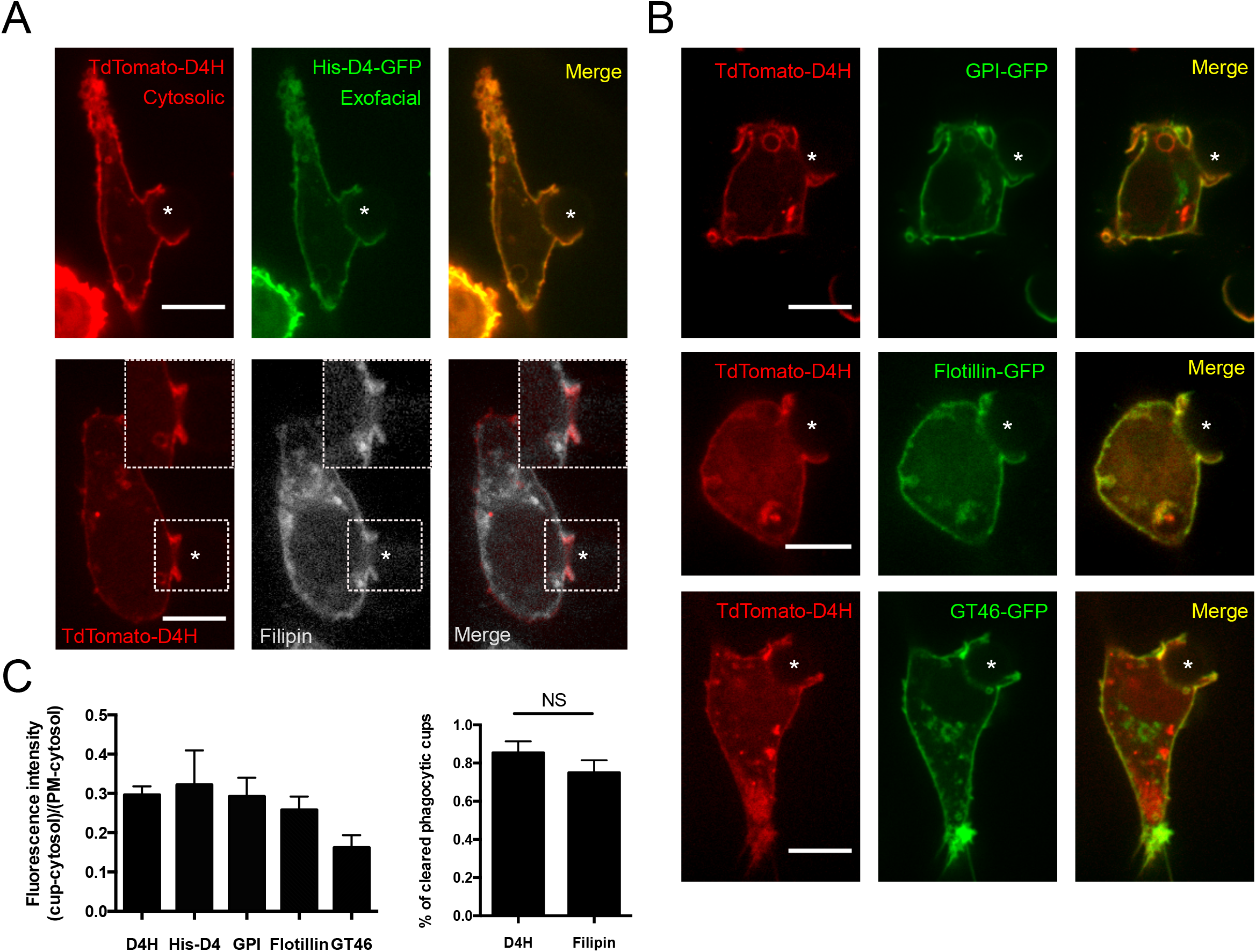
D4H is not lost at the cup due to flopping of loss of lipid raft components. **(A)** RAW macrophage co-transfected with TdTomato-D4H and labelled with 10 μg/mL His-D4H-GFP or fixed with 4% PFA and labelled with 0.5mg/mL filipin for 2 hours at 4°C. Asterisks denote phagocytic target. Insets show enlarged area. Scale bar, 10 μm. **(B)** RAW macrophage co-transfected with TdTomato-D4H and GPI-GFP, Flotillin-GFP or GT46-GFP and given IgG opsonized polystyrene beads. Asterisks denote phagocytic target. Scale bar, 10 μm. **(C)** Quantification of the fluorescence intensity of D4H and markers at the phagocytic cup (left). Data are the ratio of fluorescence intensity of D4H or markers at the phagocytic cup to the PM + SEM of three independent experiments. The equation (cup-cytosol)/(PM-cytosol) was used for calculations. Quantification of the percentage of cleared phagocytic cups (right). Data are the percentage of phagocytic cups with D4H or filipin cleared + SEM of three independent experiments (right). 5-24 cells were analyzed for each marker. A Student’s two-tailed unpaired *t*-test was performed. NS, not significant.

### Cholesterol-rich and -poor domains are both depleted during phagocytosis

Membrane rafts are thought to act as a scaffold to mediate signal transduction events and suggest a role for cholesterol during FcγR signaling^11–14^. Therefore, a reduction in cholesterol from the base of the phagocytic cup could be due to the preferential loss or displacement of cholesterol-rich membrane domains. To this end, we examined whether the membrane raft markers, flotillin and GFP-GPI, are depleted explicitly during phagocytosis. As shown in **Figure 2B and C**, flotillin and GFP-GPI display a similar localization as the D4H during the phagocytosis process. However, a similar pattern was observed for a non-membrane raft resident protein, GT46, a synthetic fusion protein comprised of the signal sequence of lactase-phlorizin, GFP, a consensus N-glycosylation site, the transmembrane domain of the LDL receptor and the cytoplasmic tail of CD46^15^. Together these results demonstrate that both raft and non-raft components are equally diminished during phagocytosis.

### Focal exocytosis of endosomes locally dilutes cholesterol

Extensive membrane remodeling occurs during phagocytosis with both focal endocytosis and exocytosis occurring during the progression of phagosome formation^16^. To test whether endocytosis and exocytosis are contributing to the reduction of cholesterol from the base of the phagocytic cup, we inhibited each process individually and monitored cholesterol using D4H. To block clathrin-mediated endocytosis, we treated macrophages with Dyngo 4a, an inhibitor of dynamin, a GTPase required for the scission of vesicles from the PM. As a control, Dyngo 4a was a potent inhibitor of transferrin endocytosis (**Figure 3A**). However, during phagocytosis of IgG-opsonized particles, Dyngo 4a had little impact on the dynamics of cholesterol as monitored by the D4H probe (**Figure 3B, C**). Thus, while endocytosis is occurring during phagosome formation, it is not removing a significant portion of the cholesterol-rich membrane.

**Figure 3.**
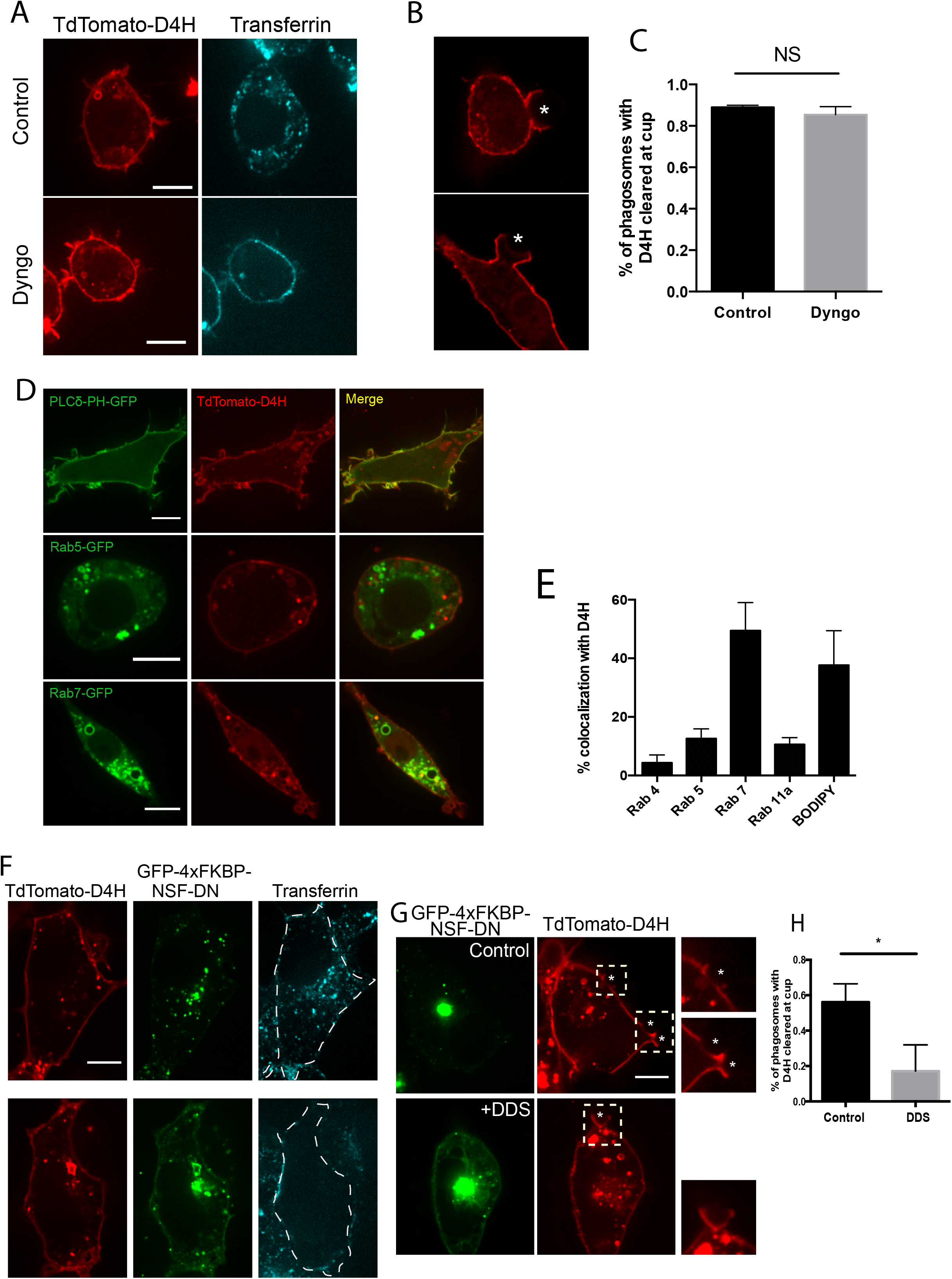
D4H is lost at the cup due to exocytosis. **(A)** RAW macrophage transfected with TdTomato-D4H and treated with 15 μM dyngo for 15 minutes at 37°C or left untreated, and 50 μg/ml transferrin for 15 minutes at 37°C. Scale bar, 10 μm **(B)** RAW macrophage transfected with TdTomato-D4H and treated with 15 μM dyngo or left untreated and given IgG opsonized polystyrene beads. Asterisks denote phagocytic target and inset shows enlarged area of the cup. Scale bar, 10 μm. **(C)** Quantification of the percentage of phagocytic cups with D4H cleared. Data are the percentage + SEM of three independent experiments with at least 10 cells analyzed per condition. A Student’s two-tailed unpaired t-test was performed. NS, not significant. **(D)** RAW macrophages co-transfected with TdTomato-D4H and PLCδ-PH-GFP, Rab5-GFP or Rab7-GFP or transfected with TdTomato-D4H and labelled with 0.01 mg/ml BODIPY at 37°C for 20 minutes in serum free medium. Scale bar, 10 μm. **(E)** Quantification of D4H colocalization with endosomal markers, percent colocalization represent the number of D4H puncta that were positive for the corresponding markers. Data represent the means of three independent experiments + SEM. At least 80 puncta were assessed per condition. **(F)** COS-IIA cells co-transfected with TdTomato-D4H and GFP-4xFKBP-NSF-DN and treated with 0.5 μM DDS for 1.5 hours or left untreated at 37°C, and 50 μg/ml transferrin for 15 minutes 37°C. Scale bar, 10 μm. **(G)** COS-IIA cells co-transfected with TdTomato-D4H and GFP-4xFKBP-NSF-DN and treated with 0.5 μM DDS for 1.5 hours or left untreated at 37°C before given IgG opsonized polystyrene beads. Asterisks denote phagocytic target and inset shows enlarged area of the cup. Scale bar, 10 μm. **(H)** Quantification of the percentage of phagosomes with D4H cleared at the phagocytic cup in control and DDS treated cells + SEM of three independent experiments with at least 25 phagosomes analyzed per condition. P-value was calculated using a Student’s two-tailed unpaired *t*-test, * P<0.05.

The exocytosis of endosomes and lysosomes is known to be critical to support phagocytosis, especially of large particles^17,18^. We theorized that the exocytosis of relatively cholesterol-poor endosomes/lysosomes could locally dilute the cholesterol in the base of the phagocytic cup. Co-expression of D4H together with markers of endosomes, lysosomes and lipid droplets demonstrated that only a fraction of the organelles had sufficiently high levels of cholesterol to be decorated with the probe. For instance, we found that of the cytoplasmic D4H puncta ≈49% were positive for Rab7 and ≈38% were positive for the lipid droplet stain BODIPY (**Figure 3D, E**). Other endosomal markers including Rab4, Rab5, and Rab11a which are markers for fast recycling, early endosomes and slow recycling, respectively, accounted for only a small fraction of the cytoplasmic D4H punta.

To determine if exocytosis was responsible for the localized depletion of cholesterol in the forming phagosome, we inhibited the activity of W-ethylmaleimide-sensitive factor (NSF), a hexameric protein that mediates the recycling of SNARE proteins required to support exocytosis^19,20^. As prolonged and even transient expression of dominant negative (DN) forms of NSF are toxic to cells we sought to use a more acute treatment. We constructed a chimera that contains DN NSF (E329Q) attached to 4 mutant FK-506 binding protein [FKBP (F36M)] domains which are conditional aggregation domains, conjugated to GFP^21^. Expression of this fusion protein results in the formation of cytoplasmic aggregates of this protein. Upon addition of an FK-506 ligand (DDS) a portion of the aggregated protein is released, or “knocked-loose”, allowing for acute impact of the DN-NSF (**Supp Fig 1A**). Unfortunately, despite repeated attempts with variations of the protocol we could not get the DDS to work with the RAW macrophage. Indeed, macrophage are known to express a significant amount of detoxification proteins and we suspect that in these experiments we did not obtain a critical concentration of the DDS compound within the RAW cells^22^.

We took advantage of the COS1 cells expressing the FcγRIIa that displays similar cholesterol dynamics during the course of phagocytosis. As expected in control conditions, transiently tranfected GFP-4xFKBP-NSF-DN was observed to be punctate in the cytosol of COS-IIA cells (**Figure 3F**). After addition of DDS, a portion of the GFP-4xFKBP-NSF-DN is cytosolic and also visible at the PM. To monitor the impact of the GFP-4xFKBP-NSF-DN we examined the uptake of transferrin, a constitutive process that requires constant uptake and recycling of the transferrin receptor and as a result is inhibited by NSF-DN. As expected in cells expressing the aggregated GFP-4xFKBP-NSF-DN, but in the absence of the solubilizer the fluorescent transferrin was readily internalized and transported to the recycling endosomes. While in cells expressing the GFP-4xFKBP-NSF-DN and treated for 1.5 hours with the solubilizer show no uptake of fluorescent transferrin and even an absence of binding to the cell surface consistent with expression of NSF-DN. Next, we examined the impact of the knocked-loose GFP-4xFKBP-NSF-DN on the cholesterol dynamics during phagocytosis. In cells treated with the DDS, GFP-4xFKBP-NSF-DN was observed in both the PM and at the phagocytic cup while in transfected cells not treated with the solubilizer, the GFP signal was mainly aggregated in the cytosol (**Figure 3G**). In these same cells we quantified the fraction of cells that displayed depleted D4H from the base of the cup. While in control COS-IIA cells nearly 60% of the cells showed a reduction of D4H, in the cells treated with the solubilizer only 20% showed considerable clearance (**Figure 3H**). This suggests that it is exocytosis and the delivery of endomembranes to the phagocytic cup that contributes to the reduction of cholesterol in the base of the phagocytic cup.

### Diminution of cholesterol and Lyn kinase displacement

Our observation that there is a localized reduction in cholesterol content at the base of the forming phagosome, together with lipidomics data demonstrating that there is less cholesterol in early phagosomes compared to the PM^5^, led us to ponder if these dynamic changes in cholesterol had any biological significance. Previous results had demonstrated that activation of the homologous FcεR results in the transition of Lyn kinase from Triton X-100 soluble to insoluble fractions^23^ and that cholesterol-rich membrane domains promoted Lyn kinase activity in mast cells^24^. Conversely, removal of cellular cholesterol decreases tyrosine phosphorylation and prevents the association of Lyn and activated FcεR with detergent resistant membranes^25,26^. Lyn kinase possesses two lipid modifications, myristoylation and S-palmitoylation, required for localization to the PM. In macrophage the association of FcγR with detergent resistant membranes is also necessary for tyrosine phosphorylation leading to the conclusion that cholesterol-rich domains are important during phagocytosis^27^. Consistent with this notion is the observation that following activation, FcγRII and Lyn are found in detergent resistant membranes while removal of cholesterol with mβCD displaces Lyn thereby preventing FcγR phosphorylation. This demonstrates that membrane rafts that are enriched with cholesterol can act to mediate Lyn kinase activity^28^.

We sought to investigate Lyn kinase signaling and tyrosine phosphorylation during FcγR-mediated phagocytosis in RAW macrophage. As shown in **Figure 4A**, the loss of cholesterol during phagocytosis is also accompanied by a nearly equivalent reduction in Lyn. However, this was not due to displacement of the phagocytic receptor as FcγRIIa remains present throughout the phagocytic cup. This observation raises the possibility that a localized depletion of cholesterol and the corresponding displacement of Lyn could help to facilitate the termination of phosphotyrosine-based signaling from the base of the cup once it is no longer needed as the pseudopods advance around the target. To look at this possibility in greater detail we switched to a two-dimensional model of phagocytosis termed “frustrated” phagocytosis and immunostained for phosphotyrosine (**Figure 4B**). As shown in the XY panel from **Figure 4B**, D4H and Lyn both clear from the centre of the frustrated cup with much of the fluorescent signal coming from organelles in the cytoplasm (XZ projection). This clearance could be detected as early as 2 min after the initiation of frustrated phagocytosis with the depleted area being maintained even after expansion of the phagocytic cup has stopped (10 min). Having shown that the D4H and Lyn are largely cleared from the PM engaging the IgG-opsonized glass coverslip, we next examined the phosphotyrosine signal. Immunostaining for tyrosine phosphorylation (pY) revealed that similar to cholesterol and Lyn kinase, the pY signal also begins to clear from the centre of the frustrated cup (**Figure 4C**). To quantify the signal intensity, we sought to use an analysis not biased by the absolute size of the cell. Thus, we measured the fluorescence intensity of the outer, middle and inner regions of the cell as depicted in the schematic in **Figure 4C** with each shell of the analysis at 33% of the frustrated cup area. Quantifying the fluorescence intensities of the shells demonstrates that there is loss of tyrosine phosphorylation within the middle of the frustrated phagocytic cup. It is worth noting that simply displacing the kinase will not be sufficient to eliminate the pY signal and that the actions of phosphatases will be required. However, the results suggest that the local depletion of Lyn kinase will limit unneeded secondary rounds of phosphorylation once this portion of the membrane is no longer actively engaged in the internalization process.

**Figure 4.**
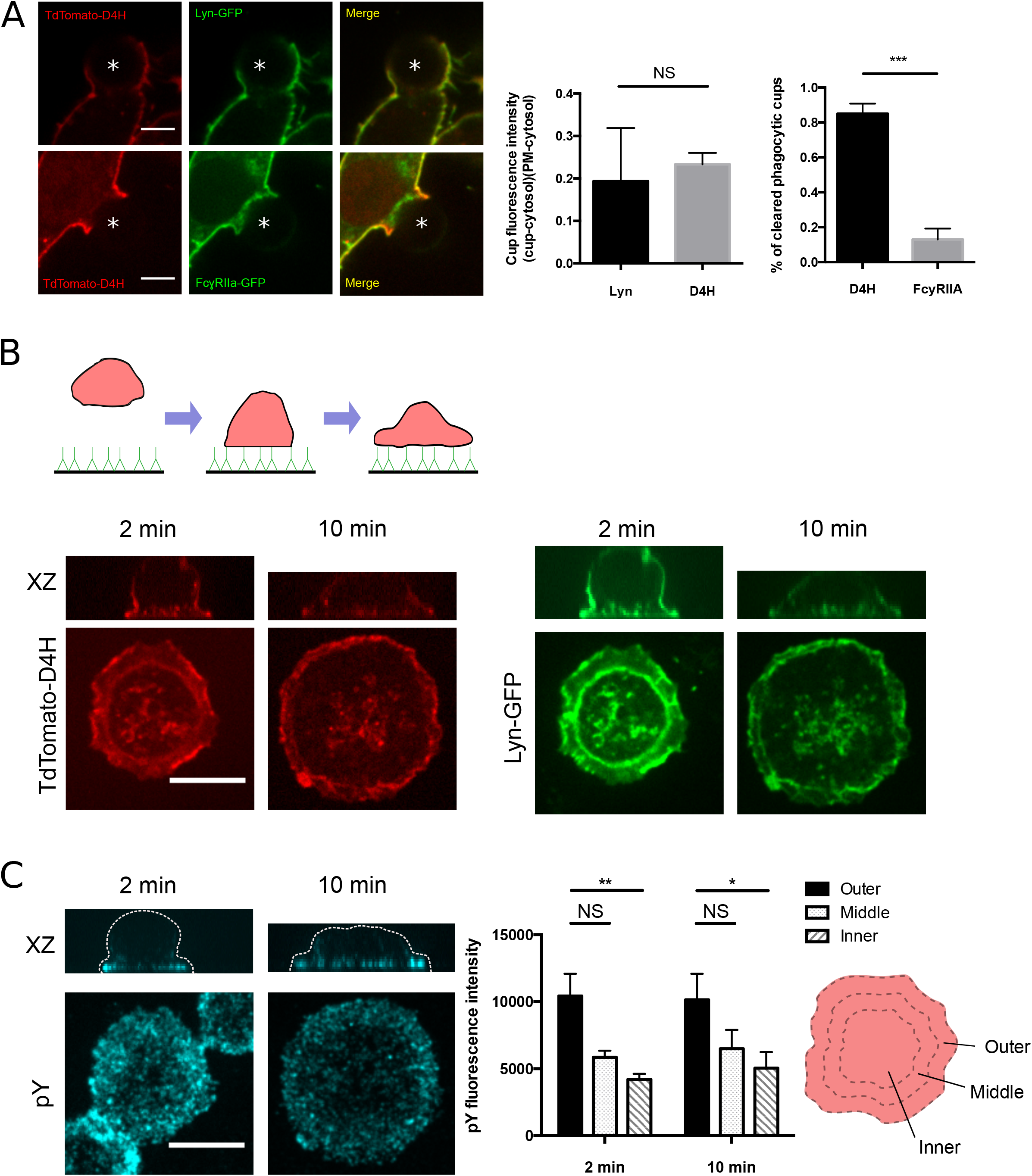
Lyn kinase and tyrosine phosphorylation terminate phagocytosis signaling. **(A)** RAW macrophage co-transfected with TdTomato-D4H and Lyn-GFP or FcγRIIa-GFP and given IgG opsonized polystyrene beads. Asterisks denote phagocytic target. Scale bar, 10 μm. Quantification of the fluorescence intensity of D4H and Lyn at the phagocytic cup (left). Data are the ratio of fluorescence intensities of D4H or Lyn at the phagocytic cup to the PM + SEM of three independent experiments with at least 30 cells analyzed per condition. The equation (cup-cytosol)/(PM-cytosol) was used for calculations. A Student’s two-tailed unpaired t-test was used performed. NS, not significant. Quantification of the percentage of phagocytic cups with D4H or FcγRIIa cleared (right). Data are the percentage of phagocytic cups with D4H or FcγRIIa cleared at the cup + SEM of three independent experiments with at least 20 cells analyzed per condition. **(B)** Schematic of the frustrated phagocytosis assay shown. Coverslips are coated with IgG and macrophages are parachuted down onto the coverslips to monitor phagocytosis. RAW macrophage co-transfected with TdTomato-D4H and Lyn-GFP and dropped onto IgG coated coverslips. Macrophages were allowed to internalize the target for the indicated time points and fixed with 4% PFA. XZ slices shown above panel. Scale bar, 10 μm. **(C)** RAW macrophage dropped onto IgG coated coverslips and allowed to internalize the target for the indicated time points. Macrophage were fixed with 4% PFA and immunostained with pY. XZ slices shown above panel, cell is outlined in white dashed line. Images are pseudo-coloured with the scale shown on the right. Quantification of the frustrated phagocytosis assay (right). Cells were outlined as shown in the schematic. The fluorescence intensity of 33% of the total area was measured with each shell. Data are the fluorescence intensities of the indicated regions + SEM of three independent experiments with at least 25 cells analyzed per condition. P-values were calculated using a two-way ANOVA test, * P<0.05, **P<0.01, NS not significant.

In summary, we characterized the dynamics of cholesterol during IgG-mediated phagocytosis using genetically encoded and recombinant versions of a cholesterol biosensor and filipin. Our results are consistent with previous findings that have demonstrated that the localized exocytosis of endosomes remodels the forming phagosomal membrane and can dilute the plasmalemmal resident proteins and lipids present at the site of phagocytosis^20^. The impact of this is two-fold. First, the exocytosis of endosomes helps to quickly remodel the membrane so that the nascent phagosome is less PM-like and more endosomal-like in nature. Second, the displacement of Lyn kinase, despite the presence of IgG-engaged FcγR helps to prevent secondary rounds of signaling once phosphatases have attenuated the signal at the base of the phagocytic cup.

## Materials and Methods

### Reagents

5.19μm and 8μm polystyrene microspheres containing 2% divinylbenzene were purchased from Bangs Laboratories, Inc. (Burlington, Ontario). Methyl-β-cyclodextrin (mβCD) was from Sigma-Aldrich (Oakville, Ontario). Cholesterol was purchased from Steraloids (Newport, Rhode Island). Dyngo 4a was purchased from Abcam (Toronto, Ontario) and DDS was purchased from Clontech (Mountain View, California). Fluorescent secondary antibodies against human IgG were from Jackson ImmunoResearch Laboratories, Inc. (West Grove, Pennsylvania). Alexa Fluor 647 conjugated transferrin and BODIPY 493/503 were purchased from Thermo Fisher Scientific (Burlington, Ontario).

### Plasmids

The plasmids used in this study were previously generated TdTomato D4H^4^, His-D4H-GFP^4^, Flotillin1-GFP^4^, FcγRIIa-GFP^6^, PLCδ-PH-GFP^29^, GFP-LactC2^30^, GPI-GFP^31^, GFP-GT46^15^, GFP-P4M-SidMx2^32^, Lyn-GFP^33^, GFP-4xFKBP-NSF-DN^34^. To generate the regulated aggregation domain plasmid EGFP-4xFKPB(F36M)-NSF(E329Q), the signal sequence from the plasmid pC_4_S_1_-ssEGFP-4xFKBP(F36M)-hGH (ARIAD, Inc.)^21,34^ was first removed. NSF (E329Q) was amplified by PCR using the pcDNA3.1-NSF (E329Q) plasmid DNA as template^19^ incorporating 5’ Spe1 and 3’ BamH1 sites. The hGH was excised from the pC_4_S_1_-EGFP-4xFKBP(F36M)-hGH plasmid with Spe1/BamH1 and the NSF (E329Q) was cloned into this site. The plasmid was sequenced to confirm the presence of the E329Q mutation in NSF. Primer sequences used to generate the constructs are available upon request.

### Cell culture and DNA transfection

The murine macrophage cell line RAW264.7 was obtained from the American Type Culture Collection (ATCC) and grown in RPMI supplemented with 10% heat-inactivated fetal bovine serum (Wisent) at 37°C under 5% CO_2_. COS-1 monkey fibroblast cell line was obtained from ATCC and grown in DMEM supplemented with 10% heat-inactivated fetal bovine serum at 37°C under 5% CO_2_. COS-1 cells stably expressing the Fcγ receptor IIA were used for experiments in phagocytosis (COS-IIA)^6^.

Macrophages were transfected with FuGENE HD (Promega) according to the manufacturer’s instructions. 100 μl of serum-free DMEM was mixed with 1 μg of plasmid DNA and 3 μl of the transfection reagent. The mix was allowed to sit for 15 minutes at room temperature before distributing equally into two wells of a 12-well plate. Cells were typically used 18-24 h post transfection. COS-IIA cells were electroporated with the Neon Transfection System (Life technologies) according to the manufacturer’s protocol. COS-IIA cells were lifted with trypsin (Wisent), counted and centrifuged at 300 *g* for 5 minutes. 5 × 10^5^ cells were resuspended in 100 μl of buffer R and incubated with 2 μg each of TdTomato-D4H and GFP-4xFKBP-NSF-DN. This solution was subjected to electroporation using two 30-millisecond pulses of 1050 V. Electroporated cells were transferred to coverslips and allowed to recover for 24 hours before experiments.

### Purification of His-D4H-GFP

His6x-GFP-D4 fusion protein was overexpressed in *E*. coli strain BL21 (Rosetta) and purified as previously described. Briefly, *E.coli* transformed with pET28b-GFP-D4 were inoculated in LB medium shaking at 37°C until OD_600_ reached 0.5. Cultures were induced with 1 mM IPTG for 4 hours at 30°C and harvested and lysed in B-PER (Pierce Biotechnology) according to the manufacturer’s instructions. Cell lysate supernatants were bound to TALON Metal Affinity Resin (Clontech) and washed with PBS. Protein was eluted with 150 mM imidazole, pH 7.57 and the fractions were ran on an SDS-PAGE gel to analyze for His6x-GFP-D4 using Coomassie Blue.

### Binding of His6x-GFP-D4 to the exofacial leaflet of the PM of cells

RAW macrophages were incubated with 10μg/mL of the recombinant protein in serum-free medium for 15 minutes at room temperature, washed with PBS and imaged live at 37°C.

### Cholesterol depletion and re-addition treatment

Sterols were dissolved in chloroform and dried with constant stream of N2 gas. Dried lipids were resuspended in mβCD/PBS and sonicated until the solution was completely dissolved. Macrophages were treated with 10 mM mβCD in serum free medium for 10 minutes at 37°C to deplete cells of cholesterol. For cholesterol re-addition, serum-starved cells were loaded with 50 μg/mL cholesterol for 30 minutes at 37°C after depleting them of cholesterol.

### Phagocytosis

RAW264.7 macrophages were plated onto 1.8 cm glass coverslips and allowed to grow for 24 hours before transfection. Polystyrene microspheres were diluted 10-fold in PBS and opsonized with human IgG (final IgG concentration 4 mg/mL) by incubating for 1 hour at 37°C. Excess IgG was removed by washing the beads with PBS. Opsonized beads were then labelled with fluorescent antibodies against human IgG for 15 minutes at room temperature. Phagocytosis was initiated by adding 15 μl of the bead suspension and sedimenting the particles by centrifugation (300 xg, 30 s). Cells were incubated at 37°C for phagocytosis to occur and fixed with 4% paraformaldehyde. Particles that were not internalized were labelled by staining with secondary antibodies against human IgG conjugated to a fluorophore that is different from the one used to label the particles.

### Endocytosis and exocytosis assay

RAW 264.7 macrophages cells were plated onto 1.8 cm glass coverslips and allowed to grow for 24 hours before transfection. Macrophages were treated with 15 μM Dyngo 4a at 37°C (Abcam) for 20 minutes in serum free DMEM before the addition of 50 μg/ml Alexa Fluor 647 conjugated transferrin at 37°C for 10 minutes in serum free DMEM. Fluorescently labelled 8μm polystyrene beads were used for phagocytosis after Dyngo treatment.

To release the aggregated NSF, RAW macrophages were plated on 1.8 cm glass coverslips and allowed to grow for 24 hours before transfection with Td-Tomato D4H and GFP-4xFKBP-NSF-DN. Macrophages were serum starved for 1 hour before treating with 0.5μM DDS at 37°C for 1.5 hours in serum free DMEM and 50μg/ml Alexa Fluor 647 conjugated transferrin at 37°C for 10 minutes in serum free DMEM. Fluorescently labelled 5μm polystyrene beads were used for phagocytosis after treatment of cells with DDS.

### Frustrated phagocytosis assay

IgG opsonized coverslips were prepared by incubating 1.8 cm glass coverslips with 1.87 mg/ml human IgG shaking overnight at 4°C. RAW264.7 macrophages were plated and allowed to grow for 24 h. Macrophages were transfected with FuGENE HD (Promega) according to the manufacturer’s instructions. 18-24 h post transfection, macrophages were serum starved, scraped and incubated at 37°C for 15 min. Cells were parachuted onto IgG coated coverslips and incubated at 37°C before fixing with 4% paraformaldehyde (PFA).

### Confocal microscopy

Images were acquired on a spinning disk confocal microscopy system (Quorum Technologies) consisting of an Axiovert 200M microscope (Carl Zeiss) with a 63X oil immersion objective (NA 1.4) and a 1.5X magnifying lens. The microscope is equipped with diode-pumped solid state lasers (440, 491, 561, 638 and 655 nm; Spectral Applied Research), motorized XY stage and a Piezo Z-focus drive. Images were acquired with a back thinned, cooled charge coupled device (CCD) camera (Hamamatsu) and Volocity software^35^. Fluorescence intensity measurements were performed with ImageJ.

### Statistics

Data presented are means and SEM of at least three independent experiments. For statistical analysis, unpaired t-tests were used.

## Acknowledgements

S.M.L. is a recipient of the Alexander Graham Bell Scholarship from the Natural Science and Engineering Research Council of Canada. G.D.F. is a recipient of a New Investigator Award from Canadian Institutes of Health Research (CIHR) and an Early Researcher Award from the Government of Ontario. This work was supported by a St. Michael’s Hospital New Investigator Start-up Fund and an NSERC Discovery Grant RGPIN-435595. We thank Sergio Grinstein (Hospital for Sick Children, Toronto, Canada) for helpful discussions.

## Abbreviation List

D4: Domain 4
mβCD: Methyl β cyclodextrin
NSF: *N*-ethylmaleimide-sensitive factor
PFA: Paraformaldehyde
PM: Plasma membrane
pY: Phosphotyrosine

---

**Supplementary Figure 1** Schematic of the knocked loose system. The conditional aggregation domains attached to DN-NSF self-aggregate without its ligand. Upon the addition of DDS, it releases the aggregates and allows for the protein to function.

